# Thought experiment: Decoding cognitive processes from the fMRI data of one individual

**DOI:** 10.1101/341594

**Authors:** Martin Wegrzyn, Joana Aust, Larissa Barnstorf, Magdalena Gippert, Mareike Harms, Antonia Hautum, Shanna Heidel, Friederike Herold, Sarah M. Hommel, Anna-Katharina Knigge, Dominik Neu, Diana Peters, Marius Schaefer, Julia Schneider, Ria Vormbrock, Sabrina M. Zimmer, Friedrich G. Woermann, Kirsten Labudda

## Abstract

Cognitive processes, such as the generation of language, can be mapped onto the brain using fMRI. These maps can in turn be used for decoding the respective processes from the brain activation patterns. Given individual variations in brain anatomy and organization, analyzes on the level of the single person are important to improve our understanding of how cognitive processes correspond to patterns of brain activity. They also allow to advance clinical applications of fMRI, because in the clinical setting making diagnoses for single cases is imperative. In the present study, we used mental imagery tasks to investigate language production, motor functions, visuo-spatial memory, face processing, and resting-state activity in a single person. Analysis methods were based on similarity metrics, including correlations between training and test data, as well as correlations with maps from the NeuroSynth meta-analysis. The goal was to make accurate predictions regarding the cognitive domain (e.g. language) and the specific content (e.g. animal names) of single 30-second blocks. Four teams used the dataset, each blinded regarding the true labels of the test data. Results showed that the similarity metrics allowed to reach the highest degrees of accuracy when predicting the cognitive domain of a block. Overall, 23 of the 25 test blocks could be correctly predicted by three of the four teams. Excluding the unspecific rest condition, up to 10 out of 20 blocks could be successfully decoded regarding their specific content. The study shows how the information contained in a single fMRI session and in each of its single blocks can allow to draw inferences about the cognitive processes an individual engaged in. Simple methods like correlations between blocks of fMRI data can serve as highly reliable approaches for cognitive decoding. We discuss the implications of our results in the context of clinical fMRI applications, with a focus on how decoding can support functional localization.

## Introduction

Paul Broca, whose work lay the foundations for the localization of cognitive functions in the brain, speculated that “the large regions of the mind correspond to the large regions of the brain” (“les grandes régions de l’esprit correspondent aux grandes régions du cerveau” in the French original) (Broca, 1861). Today, it is well established that broad cognitive domains, such as language, memory or motor functions, can be reliably mapped onto particular regions of an individual’s brain (Satterthwaite and Davatzikos, 2015). Although there is no one-to-one mapping between brain region and cognitive process (Cacioppo et al., 2007), functional localization has proven to be of direct practical use (Bunzl et al., 2010; Szaflarski et al., 2017). Functional magnetic resonance imaging (fMRI) is one non-invasive method allowing to localize brain functions with limited but nevertheless remarkable detail (Kanwisher, 2017). In the clinical context, fMRI plays an important role for planning surgery in patients with tumors or epilepsies, as it aids the understanding of which parts of the brain need to be spared in order to preserve sensory, motor or cognitive abilities (Stippich, 2015). To be useful for clinical diagnostics and prognostics, fMRI data must be interpretable on the level of the individual case (Dubois and Adolphs, 2016). Because in group studies idiosyncratic activity patterns can be obscured by averaging, the precise mapping of brain function in a single person has become a vanguard of fMRI research (Laumann et al., 2015; Huth et al., 2016; Gordon et al., 2017). These studies are important to deepen our understanding of how the brain works, because the functional organization of brains becomes more heterogeneous on a finer anatomical scale (Laumann et al., 2015; Poldrack, 2017). Also, when looking at increasingly smaller ‘regions of the mind’, such as the neural correlates of specific words instead of language in general, averaging on the group level can obscure the fine spatial information which allows to differentiate these contents in the individual brain (Huth et al., 2016). Single participant studies can also provide valuable impulses for the use of fMRI as a clinical tool. This includes the possibility to assess how stable results are within a single participant, and how much data should be collected to provide a reliable description of the individual’s functional brain organization (Laumann et al., 2015; Gordon et al., 2017). While the group average is a composite of many individuals, the activity map of the individual is likewise a composite of an underlying time course, consisting of many separate observations of brain activity while performing a task. Variability over the course of an fMRI session can be expected due to factors such as head movement, fatigue, increasing familiarity with the task and changes in cognitive strategies (McGonigle, 2012; Gorgolewski et al., 2013). The neuroradiologist’s interpretation of a single patient’s fMRI might therefore be substantially improved, if she knows how the patient’s cognitive states changed over time and how this relates to changes in brain activity patterns. This is particularly important if no overt behavior is collected during the fMRI task. For example, in a language production task, patients might be asked to produce words from categories such as “fruits” or “animals” in a pre-defined period of time (Woermann et al., 2003). Because overt articulation of words produces movement artifacts, the patients might be asked to use only internal speech. Without behavioral output from the patient, interpretation of fMRI results is limited by the uncertainty about whether the task was performed in the expected manner. A possible solution might be the decoding of fMRI data, in order to learn what the patient was thinking at each point in time. Decoding refers to an inference from brain activity patterns to the cognitive processes that accompanied them (Poldrack, 2006; Haynes and Rees, 2006). In clinical practice, decoding has proven to be highly valuable for communicating with unresponsive patients (Owen et al., 2006; Boly et al., 2007; Sorger et al., 2012). However, decoding methods are usually not being used in presurgical planning, where fMRI is used to learn how cognitive processes can be mapped onto the brain (i.e. encoding instead of decoding; Naselaris et al. (2011)). When interpreting an activity map, decoding might nevertheless be useful to better understand how the patient performed the task: Comparing different observations within-patient might allow to assess the stability of task performance during the fMRI session, while comparisons with healthy controls allow to assess if the task was performed in a prototypical way (Dubois and Adolphs, 2016). The present fMRI study aimed at decoding the domains of language, motor functions, visuo-spatial memory, face processing and task-free resting in a single individual. Each of these task domains is relevant for presurgical planning and can be used clinically in the individual patient (language (Woermann et al., 2003); motor (Håberg et al., 2004); visuo-spatial (Jokeit et al., 2001); faces (Parvizi et al., 2012)). We used four mental imagery tasks and one rest task, where the verbal instruction to engage one of the above mentioned functions was the only external input given to the participant, and the fMRI data was the only output the participant produced. In order to evaluate how well decoding works at the level of individual fMRI blocks, we first analyzed a set of training data to learn how predictions of each cognitive domain could be optimized using simple similarity metrics. Then, test blocks were decoded regarding their cognitive domains as well as their specific contents. The study was carried out as part of a graduate course in psychology at Bielefeld University, with four groups of students making predictions for the test data.

## Methods

### 2.1 Participant

Data was collected from one healthy, 25 years old, male psychology student. The participant gave written informed consent, including written informed consent to have his brain data published online. The study was approved by the ethics committee of Bielefeld University (ethics statement 2016-171).

### 2.2 Mental imagery instructions

For the four cognitive domains of language, sensory-motor skills, visuo-spatial memory and visual processing of faces, imagery instructions were adapted from the literature: For language, a semantic verbal fluency task was used, in which the participant had to generate as many words belonging to a certain superordinate class as possible (e.g. animals, fruits; Woermann et al. (2003)). To engage motor imagery, the participant was instructed to perform different sports (e.g. tennis, soccer; Owen et al. (2006)). To test visuo-spatial memory, the participant was instructed to imagine walking to different familiar locations (e.g. school, church; Jokeit et al. (2001)). To engage face processing mechanisms, the participant was asked to imagine famous or familiar faces (e.g. actors, friends; O’Craven and Kanwisher (2000)). During time periods of resting, the participant was told to engage in a state of relaxed wakefulness. The main instructions given to the participant are outlined in Supplement S1. For each task, we tried to derive predictions about the brain areas which should be active when engaging in the respective cognitive process, based on the literature. The predictions for each task are summarized in Table 1.

**Table 1.**
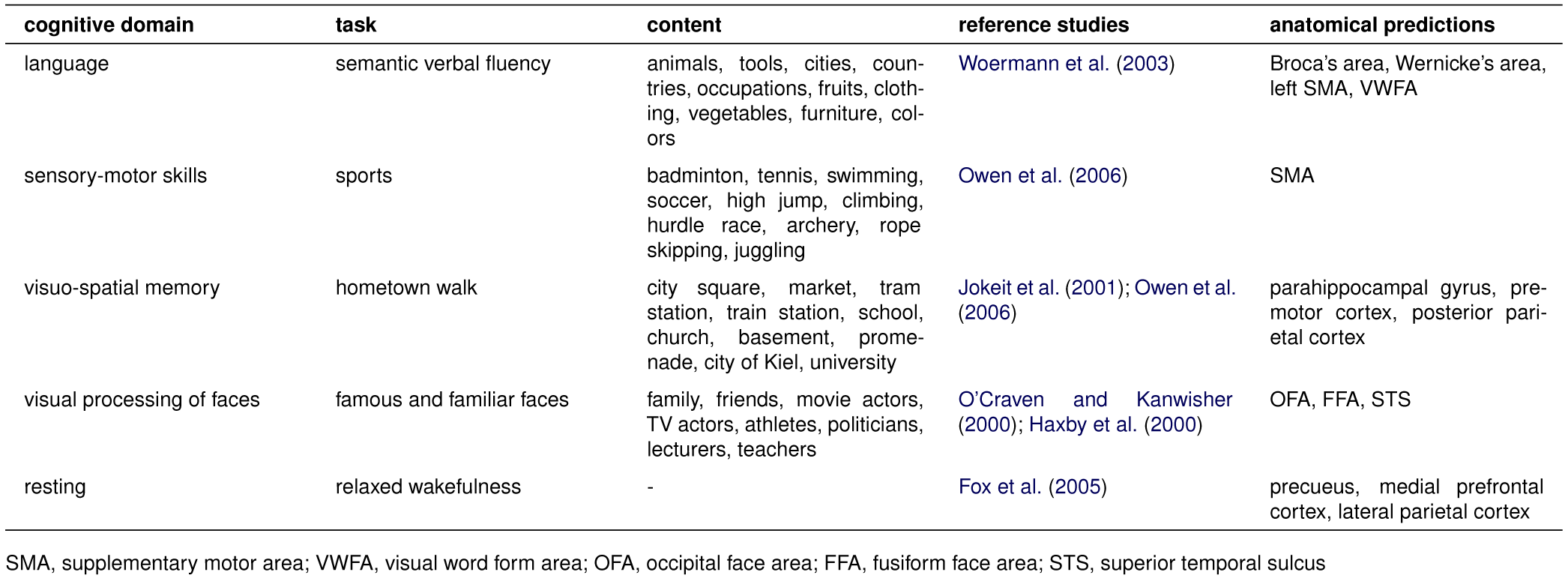
Overview of tasks used in the paradigm.

### 2.3 Study design

We acquired three runs of fMRI data, with 25 blocks per run, and a block length of 30 seconds. Within each run, there were five blocks per condition and the order of the five conditions was counterbalanced, so that they followed each other equally often. This was achieved using a simplified version of a serially balanced sequence (Nair, 1967). The full study design can be found in Supplement S2. During the experiment, the participant lay in the MRI scanner with eyes closed. Instructions to start thinking about one of the four categories and the rest condition were given by short verbal cues which were agreed upon beforehand (e.g. “language fruits”). Contents of the blocks were customized in accordance with the participant’s preferences, whenever necessary (e.g. “spatial university” will not apply to every participant’s city). Audibility was ensured by using an acquisition protocol with 1.2 second pauses between volumes, during which the instructions were given.

### 2.4 Data acquisition

MRI data were collected using a 3T Siemens Verio scanner. A high-resolution MPRAGE structural scan was acquired with 192 sagittal slices (TR=1900 msec, TE=2.5 msec, 0.8mm slice thickness, 0.75×0.75 in-plane resolution), using a 32-channel head coil. Functional echo-planar images (EPI) were acquired with 21 axial slices oriented along the rostrum and splenium of the corpus callosum (slice thickness of 5 mm, in-plane resolution 2.4×2.4 mm), using a 12-channel head coil. To allow for audible instructions during scanning, a sparse temporal sampling strategy was used (TR=3000ms with 1800ms acquisition time and 1200ms pause between acquisitions). Excluding two dummy scans, a total of 253 volumes were collected for each run. The full raw data are available on OpenNeuro (openneuro.org/datasets/ds001419).

### 2.5 Data preprocessing

Basic preprocessing was performed using SPM12 (www.fil.ion.ucl.ac.uk/spm). Functional images were motion corrected using the realign function. The structural image was co-registered to the mean image of the functional time series and then used to derive deformation maps using the segment function (Ashburner and Friston, 2005). The deformation fields were then applied to all images (structural and functional) to transform them into MNI standard space and up-sample them to 2mm isomorphic voxel size. The full normalized fMRI time courses are available online (doi.org/10.6084/ m9.figshare.5951563.v1). All further preprocessing steps were carried out using Nilearn 0.2.5 (Abraham et al., 2014) in Python 2.7. To generate an activity map for each of the 75 blocks, each voxel’s time course was z-transformed to have mean zero and standard deviation one. Time courses were detrended using a linear function and movement parameters were added as confounds. Then TRs were grouped into blocks using a simple boxcar design shifted by 2 TR (the expected shift in the hemodynamic response function) and averaged, to give one averaged image per block. These images were used for all further analyses and are available on NeuroVault (neurovault.org/collections/3467).

### 2.6 Data analysis

Emulating the “common task framework” (Liberman, 2015; Donoho, 2017), the study’s data were analyzed with regard to a clearly defined objective and a metric for evaluating success. In the “common task framework”, data for training are shared and used by different parties. The parties try to learn a prediction rule from the training data, which can be applied to a set of test data. Only after the predictions have been submitted, is the prediction of test data evaluated. It can then be explored how different approaches to prediction compared to one another, given the same dataset and objective. Accordingly, the first two fMRI runs (50 blocks total, 10 blocks per condition) of our study were used as a training set and the third fMRI run (25 blocks total, 5 blocks per condition) was used as the held-out test set. To ensure proper blinding of test data, the block order was randomly shuffled and the 25 blocks were then assigned letters from A to Y. The true labels of the blocks were only known by the first author (MW), who did not participate in making predictions for the test data. Fifteen of the authors formed four groups. Each group had to submit their predictions regarding the domain (e.g. “motor imagery”) and specific content (e.g. “tennis”) for each block in written form. The authors making the predictions were all graduate students of psychology, enrolled in a project seminar at Bielefeld University. Only after all predictions were submitted were the true labels of the test blocks revealed. The groups were allowed to analyze the training and test data in any way they deemed fit, but all used a combination of the following methods: (i) Visual inspection with dynamic varying of thresholds using a software such as Mricron or FSLView. (ii) Voxel-wise correlation of brain maps from the training and the test set, to find the blocks which are most similar to each other. (iii) Voxel-wise correlations of brain maps with maps from NeuroSynth (Yarkoni et al., 2011), to find the keywords from the NeuroSynth database whose posterior probability maps are most similar to the participant’s activity patterns. The basic principles of these analyses are presented in the following sections of the manuscript. Full code is available online (doi.org/10.5281/zenodo.1323665).

### Similarity of blocks

For similarity analyses, Pearson correlations between the voxels of two brain images were computed. This was done either by correlating the activity maps of two individual blocks with each other, or by correlating an individual block with an average of all independent blocks belonging to the same condition. During training, a nested cross-validation approach was established, where the individual blocks from one run were correlated with the averaged maps of the five conditions from the other run. Each block was then assigned to the condition of the other run’s average map it correlated strongest with. This was done for all blocks to determine the proportion of correct predictions. To learn from the training data which features allowed for the highest accuracy in predicting the domain of a block, the mask used to extract the data and the amount of smoothing were varied: Different brain masks were defined by thresholding the mean z-score maps for each of the five conditions on different levels of z-values and using only the remaining above-threshold voxel with highest values for computing correlations. The size of the smoothing kernel was also varied in a step-wise manner. The best combination of features (amount of voxels included and size of smoothing kernel used) from the cross-validation of the training data could then be used to decode the test data.

### Similarity with NeuroSynth maps

In addition to these within-participant correlations, each block was also correlated with 602 posterior probability maps derived from the NeuroSynth database (Yarkoni et al., 2011). From the 3169 maps provided with NeuroSynth 0.3.5, we first selected the 2000 maps with the most nonzero voxel. This allowed to exclude many maps for unspecific keywords such as “design” or “neuronal”, with which no specific activation patterns are associated. The selected maps were then clustered using K-Means, as implemented in Scikit-learn 0.17 (Pedregosa et al., 2011). K-Means clustering was performed starting with two clusters and then successively increasing the number of clusters to be identified. For solutions of nine or more clusters, groups of keywords representing language, auditory, spatial, motor, reward, emotion, default mode and visual processing emerged, plus additional large clusters of further unspecific keywords which were still present in the dataset (e.g. “normalization”, “anatomy”). To exclude these unspecific keywords, we eliminated the largest cluster of the nine cluster solution and re-ran the K-Means clustering on the remaining 602 maps. This clustering resulted in the same eight interpretable clusters found previously (Fig 1). To visualize the similarity between the clusters and the relationship of keywords within each cluster, we computed the Euclidean distances between all maps and projected the distances into two dimensions using multi-dimensional scaling (MDS; cf. Kriegeskorte et al. (2008)) as implemented in Scikit-learn. The resulting ’keyword space’ showed a strong agreement between the clustering and MDS, with keywords from the same cluster being close together in space (Fig 1). This ’keyword space’ was then used for decoding, by correlating our fMRI data with all NeuroSynth maps. The resulting correlations were then visualized in the 2D space, allowing to inspect not only which keywords correlated the strongest, but also if there were consistent correlations within each cluster. To be computationally feasible, a gray matter mask with 4×4×4mm resolution was used for computing correlations, reducing the number of voxel to be correlated from ~230,000 to ~19,000.

**Figure 1.**
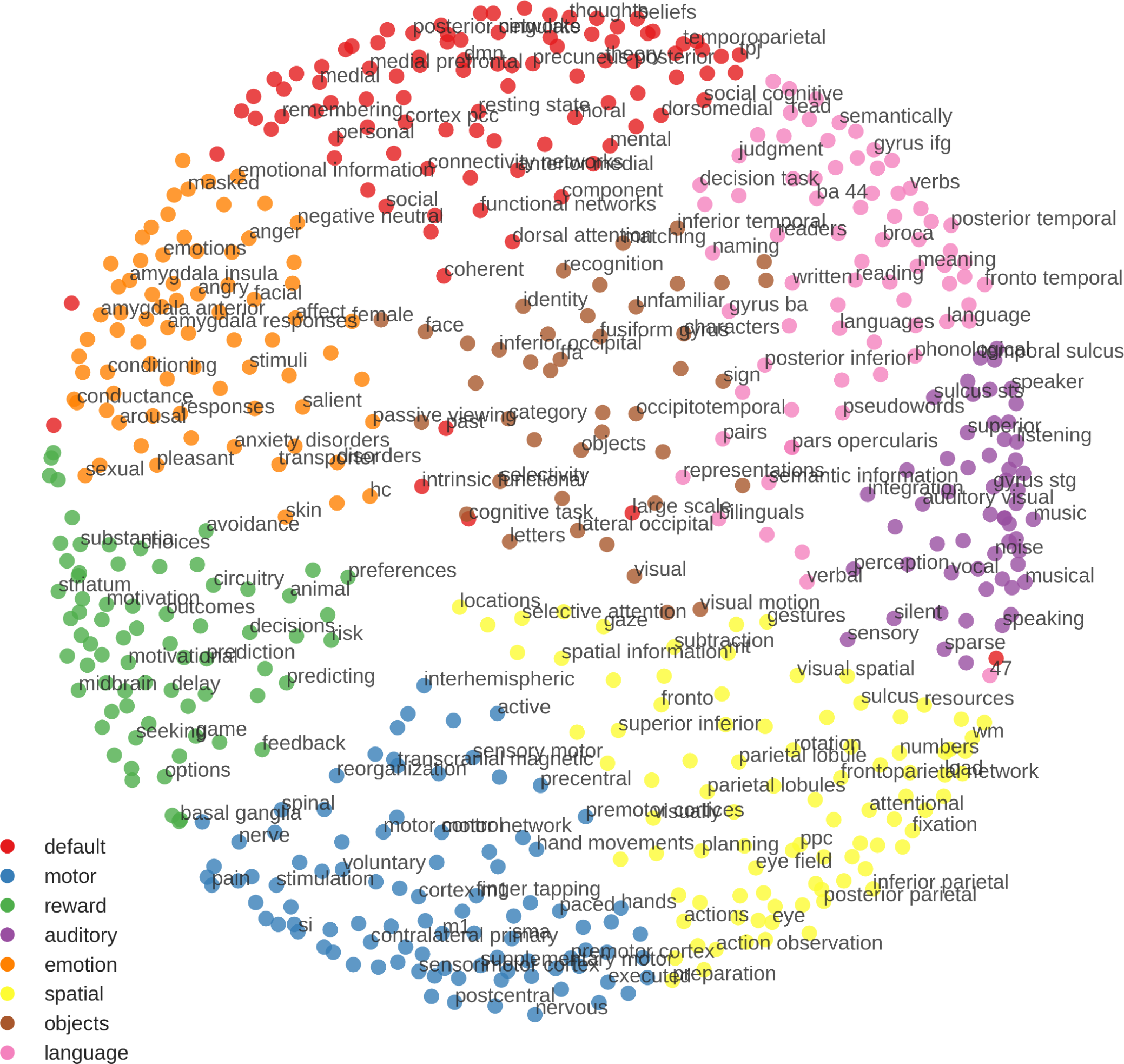
Keyword Space derived from the NeuroSynth database. Colors were assigned based on K-means clustering and distances in space were derived using multidimensional scaling (MDS). Note how both approaches give very similar results, in terms of similar colors being close together in space. There are some exceptions, i.e. BA 47 being in the default mode cluster but closer to the auditory-related keywords in MDS-space. There are clear gaps between many of the clusters, indicating that they might be categorically distinct. Regarding the arrangement of clusters, the emotion and reward clusters are close together, as well as the motor and spatial, and the language and auditory clusters. The keywords on the borders of the clusters often represent concepts shared by multiple domains, for example “characters” bridging the clusters of vision and language, “visual motion” close to vision and spatial processing, or “avoidance” related to emotion and reward processing. To allow for good readability, keywords in the figure had to be a certain distance from each other in the space to be plotted.

## Results

### 3.1 Results for the training data

#### Mean activity maps

A visualization of the average activity map for each condition is shown in Fig 2. For the language task, a clear left-lateralized network of regions, including inferior frontal gyrus, superior temporal sulcus, left supplementary motor area (SMA) and left fusiform gyrus, emerged. For the motor imagination task, SMA and premotor areas, as well as superior parietal cortex were active. The visuo-spatial memory task gave rise to activity in parahippocampal gyrus, premotor cortex and posterior parietal cortex. The face imagery condition showed activity around the mid-fusiform sulcus in both hemispheres, but mainly activity in the precuneus and medial frontal areas. For the rest condition, there was only weak activity in the precuneus, as compared to the other four conditions. Deactivations for resting were strongest in dorsolateral frontal and superior parietal regions.

**Figure 2.**
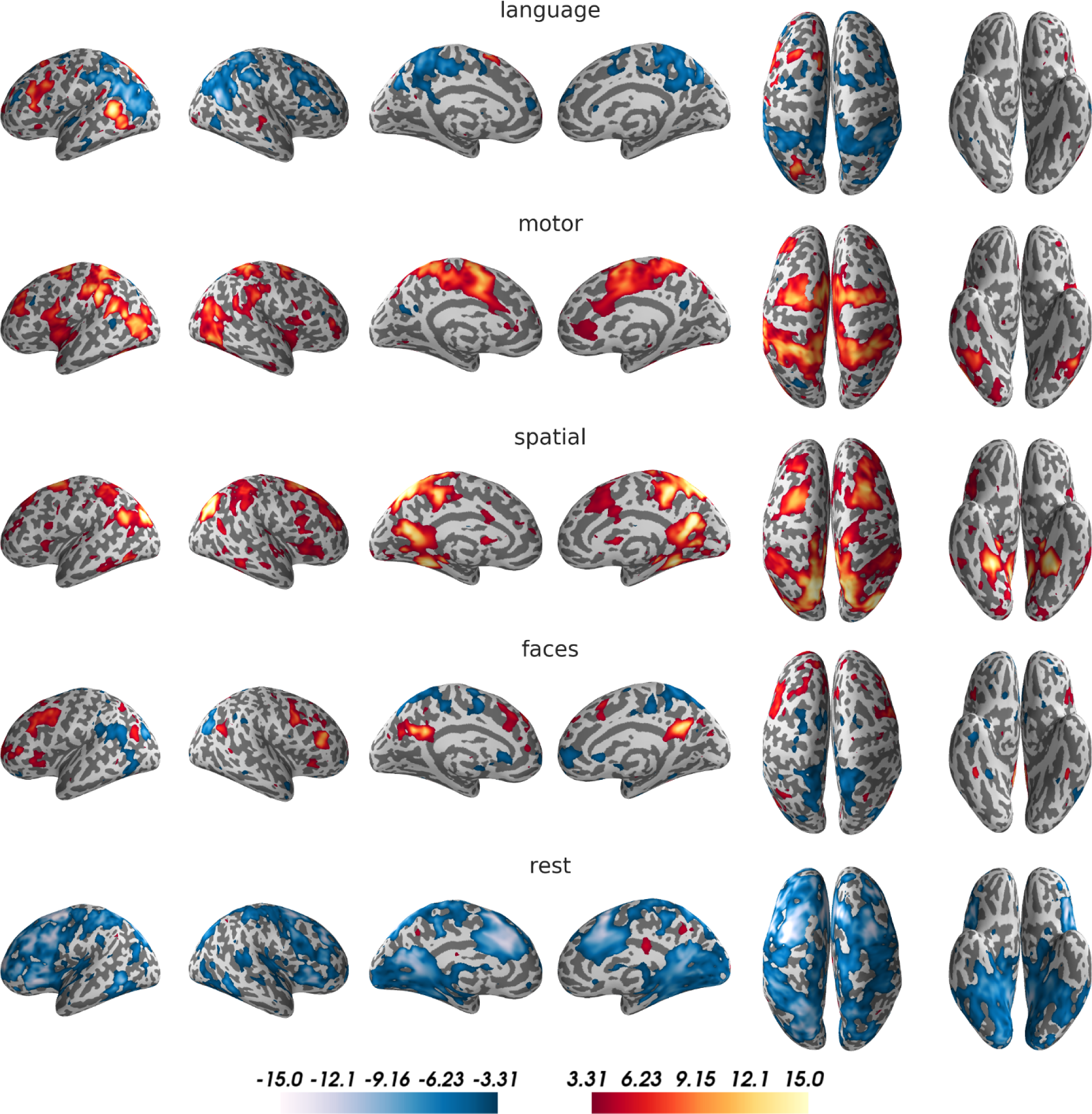
Activity maps for the five conditions of the training data. For visualization purposes, t-maps for the comparison of each condition against the remaining four were generated (smoothed with an 8mm kernel and thresholded at t=3.31, corresponding to p<0.001). Results were projected on an inflated surface of the participant’s normalized structural scan, using PySurfer. Interactive unthresholded versions of these maps are available on NeuroVault (neurovault.org/collections/3467/).

#### Feature selection for correlation analyses

For computation of similarity metrics, correlations of individual blocks from one run with the mean activity maps from the respective other run were used (i.e. blocks of run 1 correlated with the five mean activity maps from run 2, or the other way around). The decision to which domain a block belonged was then made by assigning the block to the domain it had the highest correlation with. Using this approach without voxel selection or smoothing, an accuracy of 72% was reached (p<10^-14^; for a binomial test with chance at 20%). Using feature selection (varying the voxels included and the smoothing kernel used), accuracies of up to 92% (p<10^-27^) could be reached, using only the top 1-3% of voxel from each domain and moderate or no smoothing (Fig 3).

**Figure 3.**
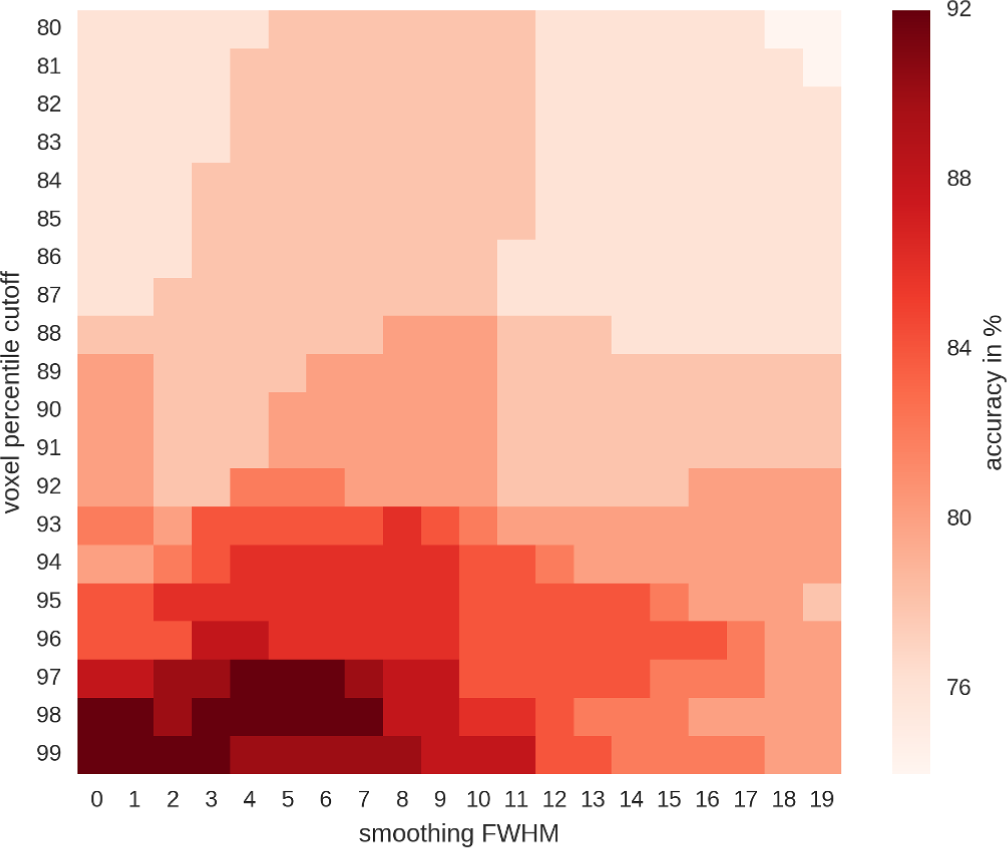
Accuracies for the predictions of training data, as a function of voxel selection and smoothing kernel. Highest accuracies (in dark red) were reached using only the top 1-3% of voxel active for each condition (i.e. using the 99-97th percentile to threshold the data). The percentile cutoff was applied to each map of the five conditions individually and the maps were then combined (conjunction of maps). Therefore, given that overlap between maps was low, percentile 80 contained 71% of whole-brain voxel and percentile 99 contained 5% of the whole-brain voxel.

The correlations of individual blocks from one run with the mean activity blocks from the respective other run, using the best feature combination, are shown in Fig 4. Of the 50 blocks, one motor block was mistaken for rest, another motor block was mistaken for a visuo-spatial memory block, and two visuo-spatial blocks were mistaken for motor blocks, corresponding to 46 out of 50, or 92%, correct predictions.

**Figure 4.**
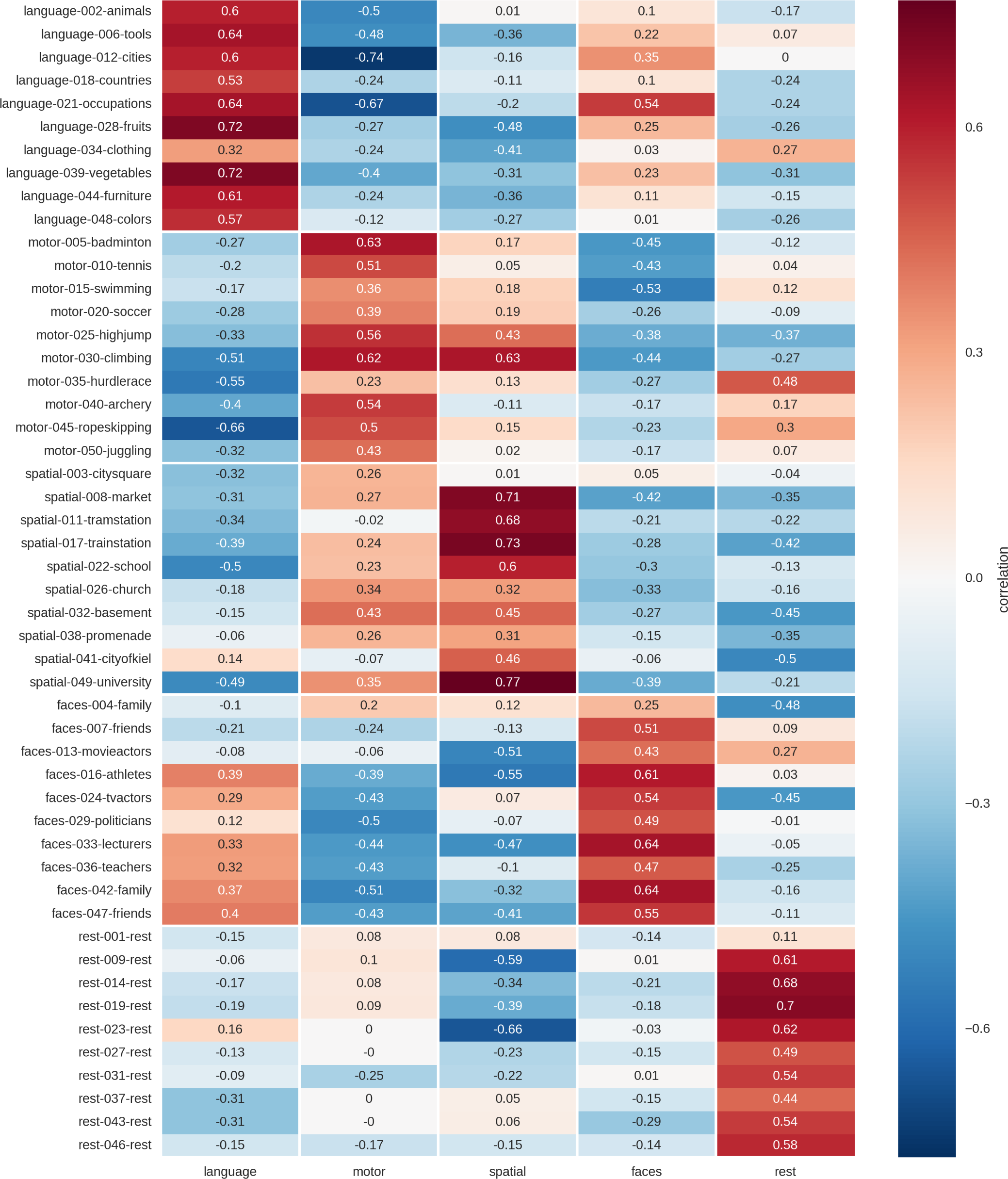
Correlation of single blocks (rows) of one run with the mean activity maps (columns) of the respective other run. Results are based on unsmoothed data using a 99th percentile cutoff to threshold the mean activity maps with which the individual blocks are correlated. For each block, the name of the condition (i.e. “language”), the number of the block in the experiment (i.e. “002” for the second block of the experiment) and the content (i.e. “animals”) are indicated in the row labels.

#### Decoding using NeuroSynth data

To evaluate how completely independent data can be used to decode the five conditions, each mean activity map (averaged over both training runs) was correlated with the NeuroSynth data and the strength of the correlation visualized in MDS space (Fig 5). Four of the five conditions showed strongest correlations with keywords from the respective related cluster (language-”reading”, motor-”motor”, visuo-spatial “spatial”, rest “theory [of] mind”). The correlations of the face condition indicated that the cognitive processes our participant engaged in during this task had more to do with episodic and working memory than with object and face processing (cf. Fig 5).

**Figure 5.**
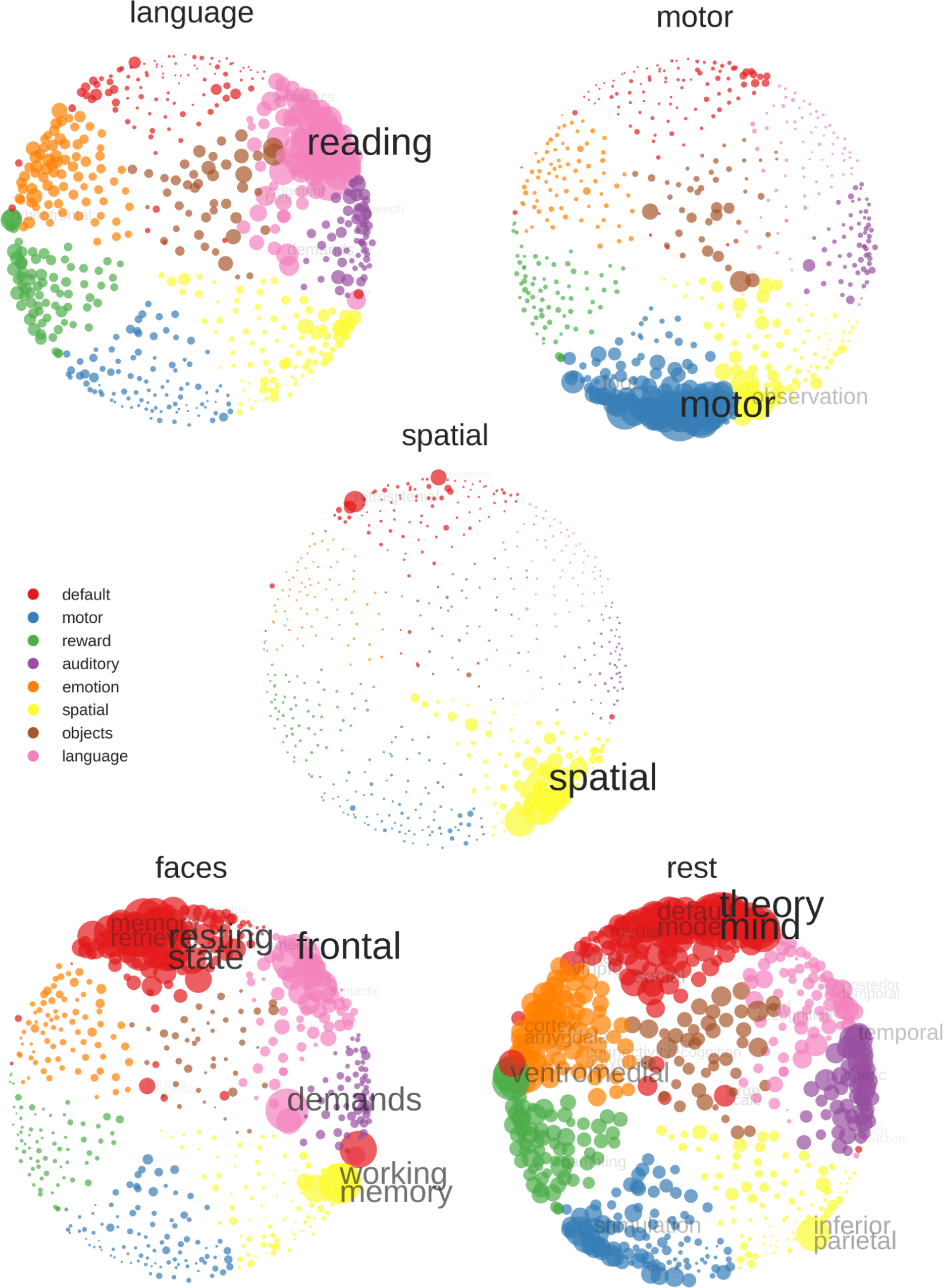
NeuroSynth decoding of average activity map for each training condition (averaged over both training runs). Stronger correlations with a keyword are indicated by a bigger circle, bigger font size and less transparency of font. To improve readability, the correlations are min-max scaled, so that the largest correlation is always of the same pre-defined size. Furthermore, the sizes of the scaled correlations have been multiplied with an exponential function, so that large correlations appear larger and small correlations smaller than they actually are (sizes are more extreme that the underlying data). To further enhance readability, if two keywords were too close in space so they would overlap, only the higher correlating keyword was printed. Color assignment is based on K-means clustering of the NeuroSynth data.

### 3.2 Results for the test set

#### Activity maps for individual blocks

Fig 6 shows the activity maps for all 25 individual blocks of the held-out test data. Here, robust activity in the networks already identified in the average training data (Fig 2) can be seen on a block-by-block basis. With the exception of block #60 for the motor imagery task, block #57 for the rest condition and block #64 for faces, specific activity in at least one of the most important regions for each domain could be found (language: superior temporal areas; motor: superior parietal areas; visuo-spatial: parahippocampal gyrus; faces: mid-fusiform sulcus; rest: precuneus).

**Figure 6.**
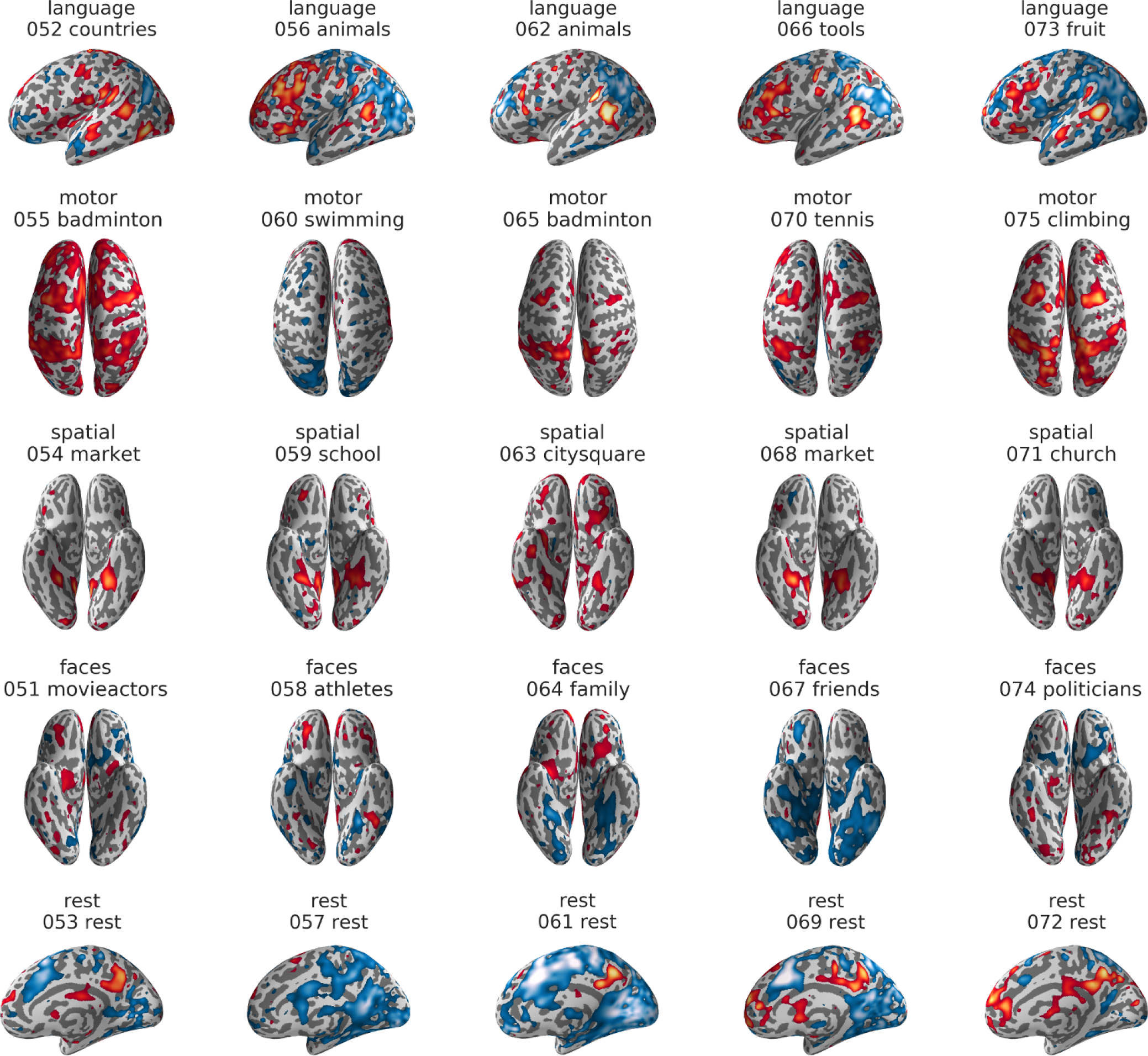
Example views of the individual activity maps of the test set. Only one view per block is shown. Maps depict the average z-values of each block, smoothed with an 8mm kernel and individually thresholded at different levels to best visualize the typical activity patterns. Red-yellow colors indicate activations and blue-lightblue colors indicate deactivations, in relation to the voxel’s grand mean over the whole timecourse. Unthresholded and interactively explorable maps of each block are available on NeuroVault (neurovault.org/collections/3467/). For each block, the name of the condition (i.e. “language”), the number of the block in the experiment (i.e. “052” for the second block of the test run, which comprises blocks 51-75) and the content (i.e. “countries”) is indicated above the brain map.

#### Correlation analysis with winner-take-all decision rule

A correlation approach using the same parameters as for the training data (top 1% of voxel, no smoothing), allowed to correctly label 24 of the 25 test blocks (96% correct; p<10^-15^; cf. Fig 7). The only misclassification occurs for block #60, where the “swimming” block from the movements condition is misclassified as belonging to the rest condition.

**Figure 7.**
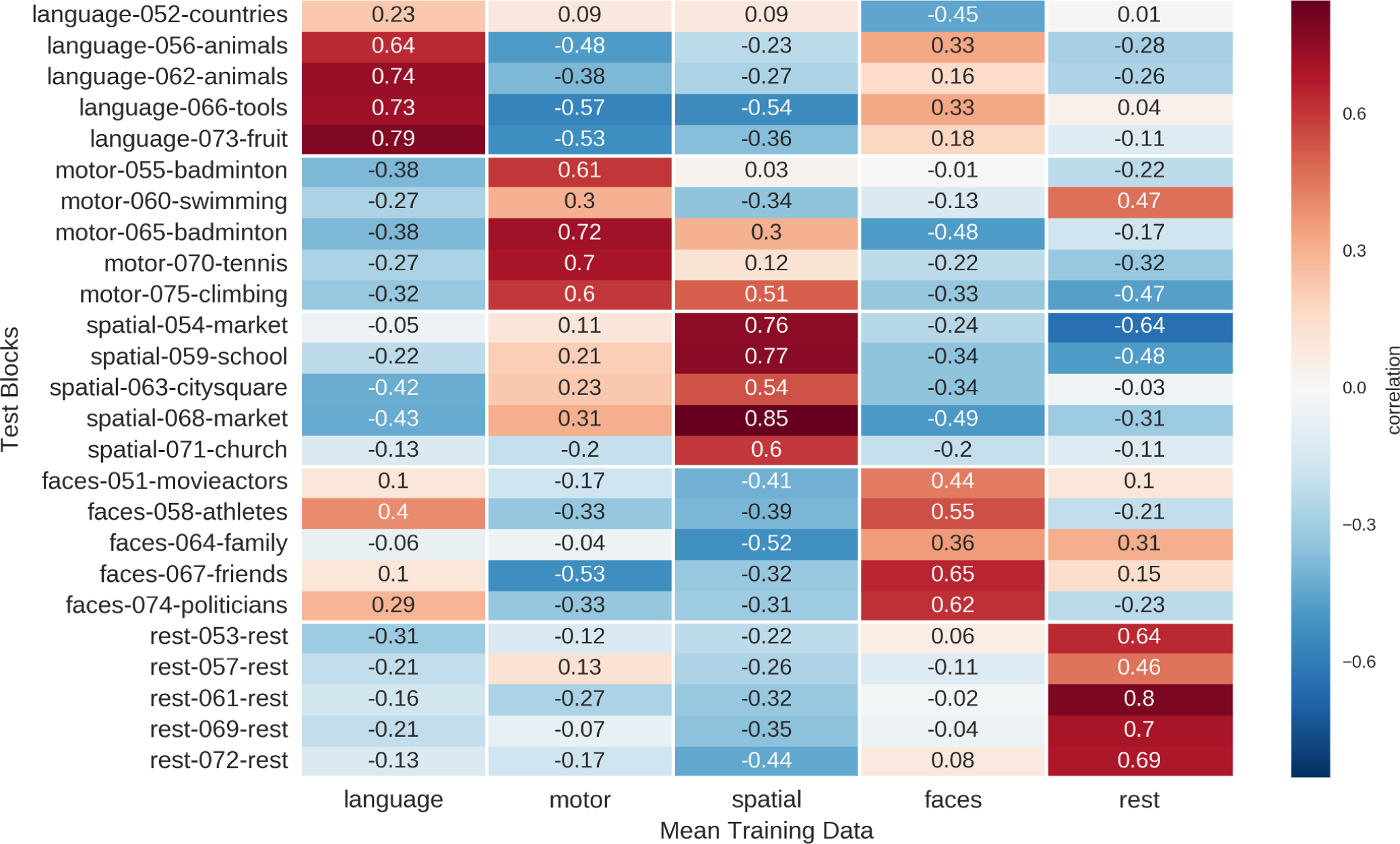
Correlation of single blocks (rows) of the test run with the mean activity maps (columns) of the two training runs. Results are based on unsmoothed data using a 99th percentile cutoff to threshold the mean activity maps with which the individual blocks are correlated. For each block, the name of the condition (i.e. “language”), the number of the block in the experiment (i.e. “052” for the second block of the test run, which comprises blocks 51-75) and the content (i.e. “countries”) is indicated in the row labels.

In addition to the correlation with mean training data, each of the 25 test blocks was also correlated with each of the 50 individual training blocks (Fig 8). Here, an optimal outcome would be if each test block had its ten highest correlations with the corresponding ten training blocks of the same condition. The results showed that for 20 of the 25 test blocks, at least eight of the highest correlations were with the correct corresponding training blocks. Only the “swimming” block had less than half of the ten highest correlations with training blocks from its correct domain.

**Figure 8.**
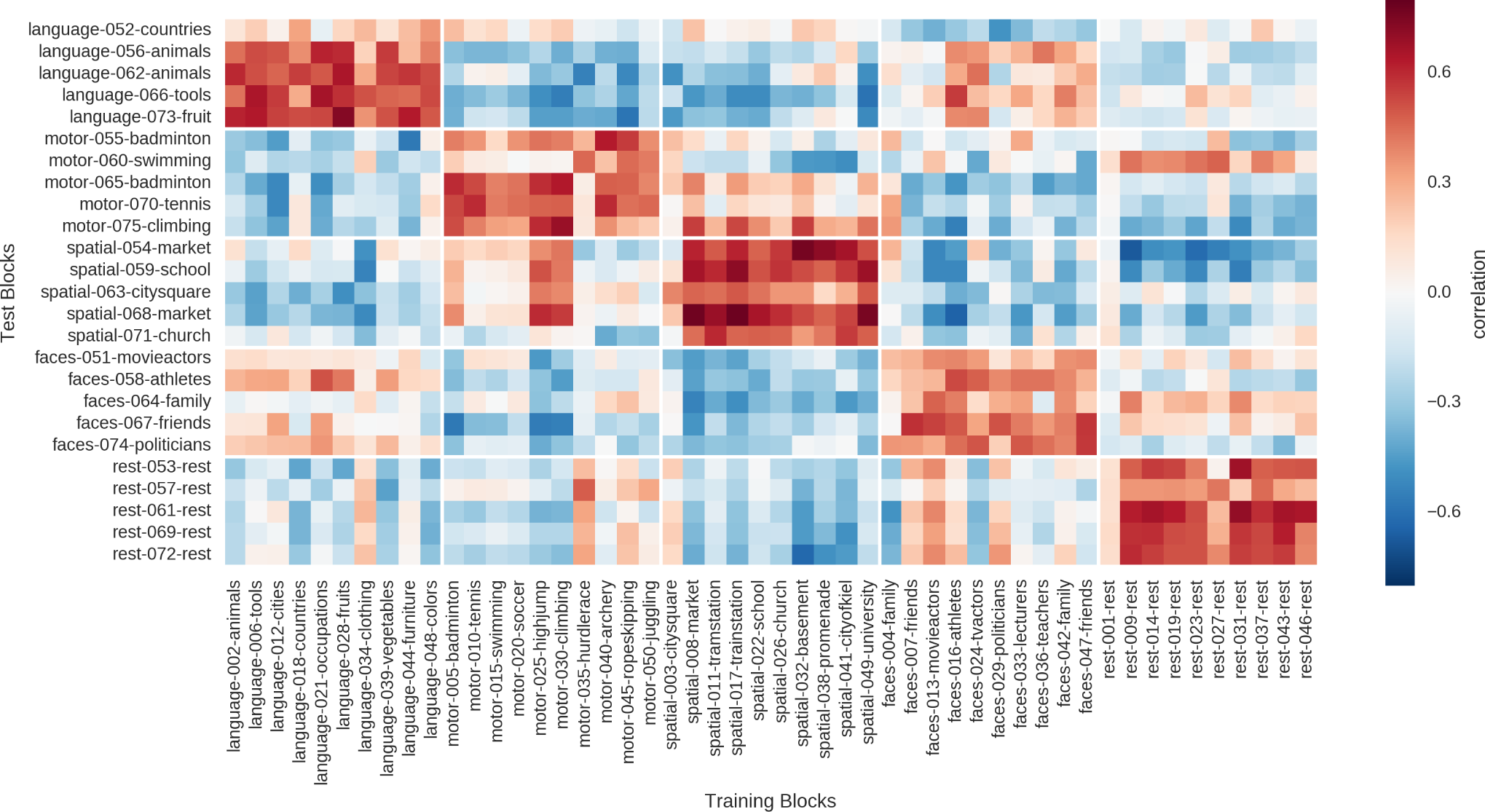
Correlation of the 25 blocks of the test run (051-075, rows) with the single blocks of the two training runs (001-050, columns).

#### Decoding using NeuroSynth data

When using the NeuroSynth data to decode each test block, 15 of the 25 of blocks (60%) were correctly decoded using the cluster of the keyword with the highest correlation (p<0.0001). The best predictions were possible for the motor, spatial and rest domains, while language and faces showed more ambiguous correlation patterns (Fig 9).

**Figure 9.**
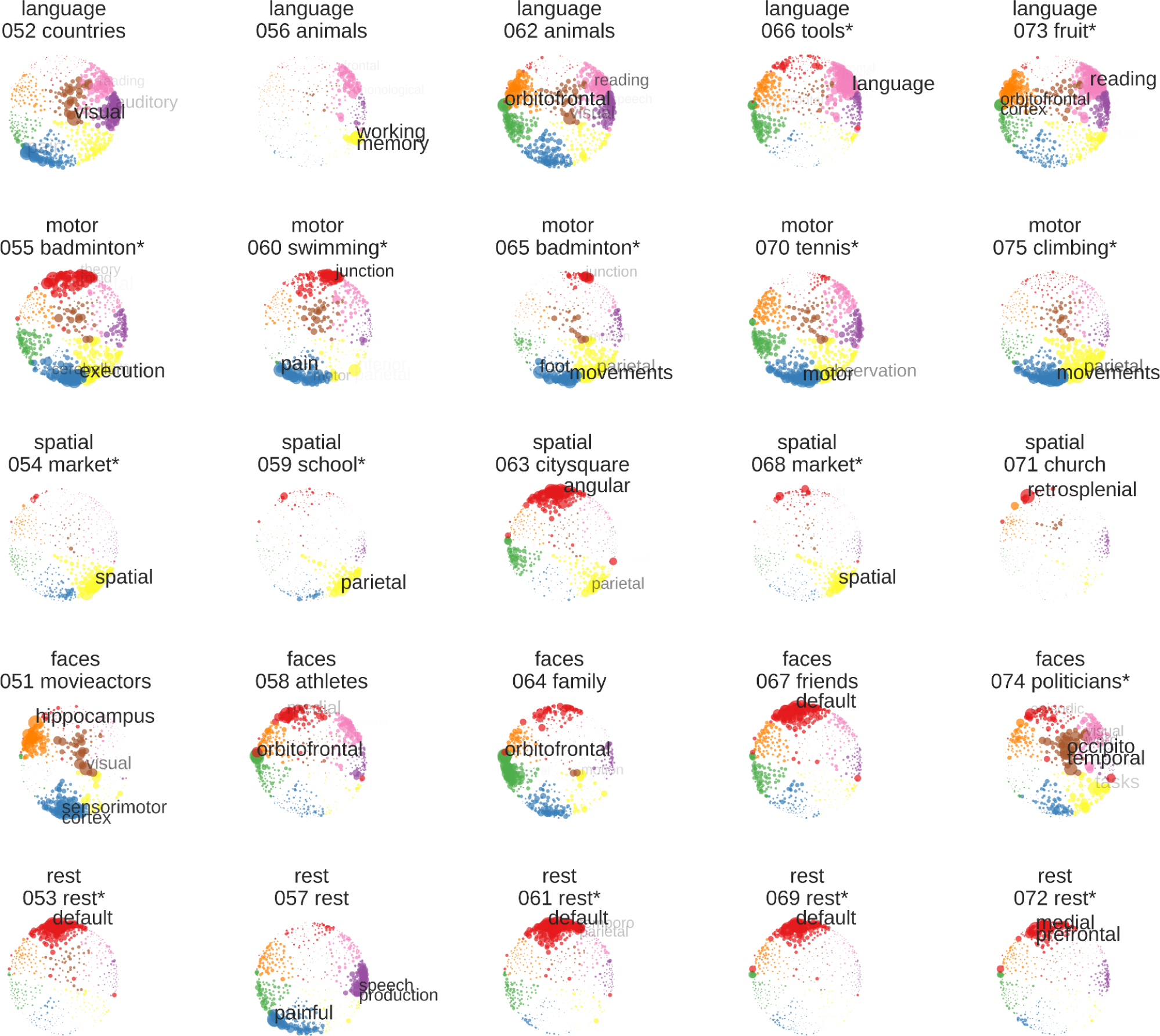
NeuroSynth decoding of individual blocks of the test run. For each block, the name of the condition, the number of the block and its content are indicated above the respective image of the space. An asterisk in the title indicates that the block was correctly decoded by assigning it to the cluster of the NeuroSynth keyword it correlated strongest with. For visualization, stronger correlations with a keyword are indicated by a bigger circle, bigger font size and less transparency of font. To improve readability, only the keywords with the highest correlations are labeled. Color assignment is based on K-means clustering of the NeuroSynth data.

#### Predictions made by the four teams

Based on these sources of information (visual inspection; correlation with mean training data; correlation with individual training blocks; correlation with NeuroSynth maps) the four teams submitted their predictions (Table 2). Three of the four teams made 23 correct predictions (p<10^-13^), all making the same mistake of classifying the swimming block as rest. In addition, the teams made at least one additional mistake, and therefore one mistake more than the correlation analysis in Fig 7. One team which weighted the results of visual inspection more strongly in their results reached an accuracy of only 76% (p<10^-8^).

**Table 2.** Results of the predictions made for the held-out test data.

| cognitive domain | correlations | NeuroSynth | group |  |  |  |
| --- | --- | --- | --- | --- | --- | --- |
|  |  |  | MH,MG,DN | SH,FH,LB,AH,SH | RV,AK,JS | DP,MS,JA,SZ |
| language | 5 (0) | 2 (-) | 5 (1)<br>fruits | 5 (3)<br>animals, tools, fruits | 5 (3)<br>animals, tools, fruits | 5 (3)<br>animals, tools, fruits |
| sensory-motor skills | 5 (2)<br>tennis, climbing | 5 (-) | 4 (1)<br>climbing | 4 (3)<br>badminton, tennis, climbing | 3 (2)<br>tennis, climbing | 4 (3)<br>badminton, tennis, climbing |
| visuo-spatial memory | 5 (0) | 3 (-) | 5 (0) | 5 (1)<br>market | 5 (0) | 5 (1)<br>market |
| visual processing of faces | 5 (2)<br>athletes, friends | 1 (-) | 5 (1)<br>movie actors | 4 (3)<br>movie actors, athletes, friends | 3 (1)<br>athletes | 5 (2)<br>athletes, friends |
| resting | 4 (-) | 4 (-) | 4 (-) | 5 (-) | 3 (-) | 4 (-) |
| total | 24 (4) | 15 (-) | 23 (3) | 23 (10) | 19 (6) | 23 (9) |

Number of correct predictions for each domain, for the 5 blocks of the test data per condition. Predictions are based on the correlation approach (top 1% of voxel, no smoothing), the NeuroSynth analysis, and based on the submissions made by the four teams of human raters. Numbers in brackets indicate the number of correct predictions regarding content. (-) indicates that content predictions were not performed. Content predictions were made for all cognitive domains except rest. Abbreviations indicate author names.

Regarding the prediction of content, the rest blocks had to be excluded, as they had no content, leaving 20 blocks from four conditions. Making the conservative assumptions that one can predict all categories perfectly and that there are only 5 possible contents within each condition, guessing would be at 20% and at least 40% correct would be needed to reach above-chance (p<0.05) accuracies. Only two of the four teams scored better than chance, with one team making 10 correct predictions out of 20 (p=0.003) and the other team 9 out of 20 (p=0.01; cf. Table 2). As all teams used a combination of all methods to guess the content, the results do not allow to infer the role each individual method played for reaching these accuracies. However, using only the automated procedure of selecting the content of the training block with the highest correlation (cf. Fig 8), only chance performance (4 out of 20) could be reached.

## Discussion

We showed that decoding “large regions of the mind” (Broca, 1861), namely language, motor functions, visuo-spatial memory, face processing, and task-free resting is possible using individual 30-second blocks of fMRI data. As in previous studies (Boly et al., 2007; Sorger et al., 2012), we were able to reach almost perfect accuracies when deciding between these cognitive domains using training data from the same participant. This was confirmed by visual inspection of the data (Fig 6), which showed that activity patterns on single block level were highly robust. We were also able to show how single blocks of a person’s fMRI data can be decoded, regarding their cognitive domains, using an independent database which maps activations of hundreds of keywords onto the brain (Yarkoni et al., 2011). This demonstrated the potential to decode a completely new observation of brain activity in a person, even when no training data were available and no feature selection had been performed. Our results also showed that it is possible to predict the contents within some of the domains with moderate accuracy. Predictions worked best for the language and motor domains, in line with previous work: The contents of our language task can be considered superordinate categories in their own right (i.e. animals and tools as animate and inanimate objects), and thus their differential activity patterns can be expected to differ on a relatively large anatomical scale (Mummery et al., 1998). The different sports used in the motor task activated different parts of the body, and could potentially be identified based on the somatotopic organization of SMA and superior parietal cortex (Fontaine et al., 2002; Aflalo et al., 2015). Because predictions of content were not explicitly trained and were only at chance using the automatic methods, the interpretation of this part of the results is limited. On the level of the five cognitive domains, the activity patterns were broadly in line with our a priori predictions (cf. Table 1). However, the motor imagery task recruited predominantly superior parietal areas, which have previously been shown to be important for movement planning (Aflalo et al., 2015), but are not always active in imagery tasks (for example Owen et al. (2006)). This also reflects the issue that while mental imagery tasks are easy to setup and integrate into the clinical routine, they have some natural limits regarding the localization of functions. In the case of the motor imagery task, one cannot reliably map the primary motor cortex (Dechent et al., 2004), where the execution of actual movements would be represented. While SMA and superior parietal areas are certainly also important for carrying out movements (Fontaine et al., 2002; Aflalo et al., 2015), it would thus be a mistake to use motor imagery as the only functional localizer in this domain. Despite this limitation of our paradigm, it is also conceivable that activity maps based on imagining complex movements could be a useful complement to simple real movement tasks, such as finger tapping. This is especially true since the organization of primary motor areas can be well approximated from brain structure alone, while this is not the case for movement planning (Aflalo et al., 2015). In contrast to our prediction, the face imagery task predominantly recruited the precuneus instead of the core face processing areas (Haxby et al., 2000). This might reflect a strong involvement of autobiographical memory recall when thinking of known faces (Gobbini and Haxby, 2007). Although unexpected, these patterns were very stable across blocks and sufficiently different from the resting activity to allow for perfect accuracies when predicting the face blocks. The resting condition produced only weak activity in the precuneus, but strong deactivations in the task-positive network, which is anti-correlated with the default mode network (Fox et al., 2005). This information was probably most important for allowing successful prediction of the rest blocks. Finally, while the activity in the superior temporal sulcus in the verbal fluency task might correspond to part of Wernicke’s area (Price, 2011), a more prototypical activity pattern would have included posterior parts of the superior temporal gyrus as well (Tremblay and Dick, 2016). It is also rather atypical that the peak of activity in a language production task is in temporal and not in inferior frontal areas (Woermann et al., 2003). Apart from that, the language and visuo-spatial conditions produced activity patterns that were very close to what would be expected if the tasks would be actually carried out, which makes these paradigms especially useful for clinical applications (Woermann et al., 2003; Jokeit et al., 2001). Another limitation of the present mental imagery task concerns the question how well activity patterns are comparable between individuals. When using external stimulation, every participant receives the same low-level inputs, but between-participant variance is still sizable (Haxby et al., 2011). Therefore, when using potentially idiosyncratic mental imagery, even larger variation between individuals should be expected. Given that we collected data from only one person, the generalization of our results is particularly limited. However, we were able to make reasonable predictions about our participant’s cognitive processes using independent data from NeuroSynth. The NeuroSynth data represents a different metric (posterior probabilities; cf. Yarkoni et al. (2011)), from different participants who performed different tasks on different scanners and were analyzed using different software and statistical methods. That our data still converged rather well with this meta-analytical information provides tentative support that the activity patterns we found were not merely idiosyncratic but to a substantial degree prototypical for the cognitive domains of interest. Furthermore, because no kind of training was performed to optimize the performance of the NeuroSynth approach, it might serve as a demonstration of ‘ad hoc’ decoding. This immediacy of application might make it especially appealing in the clinical context, where there might be no time to collect and analyze training data for each patient. However, more sophisticated methods which use the NeuroSynth database for decoding also exist (Rubin et al., 2017), and should be compared to the current approach in future studies. A crucial question regarding how the present results can inform clinical applications, is how well the present results can generalize from healthy participants to patients: While the block-wise analyses worked very well with a cognitively unimpaired and highly motivated participant, cognitive deficits, medication and a general tendency for increased movement artifacts (Van Dijk et al., 2012) will all contribute to altered or weaker signal in patient populations. Being able to collect healthy normative samples (Dubois and Adolphs, 2016) is one of the major advantages of fMRI over other methods used in presurgical planning (i.e. intracrianal EEG, Wada-testing). However, this is moot if the clinical data of actual patients cannot be reasonably collected and analyzed in the first place. Therefore, future studies are needed to show if the signal yield necessary for block-wise analyses is attainable when examining presurgical patients. It is also important to note that the four teams making predictions were very homogeneous regarding their background and approach. This leaves open the question how well the current approach would have fared against the visual inspection done by trained neuroradiologists. Also, the way in which the teams combined information from the different analyses was not made explicit. Ideally, each team would have submitted clearly formalized algorithms, which could have been compared against each other in more detail. While we outlined some important limitations above, we believe that the current study provides some valuable impulses for the clinical application of task-fMRI, including its use in presurgical planning:

### Analysis of fMRI data can benefit from splitting a dataset into smaller subsets

If the patient has consistently preformed the task as required, splitting the fMRI run into smaller parts (ideally blocks) can increase the neuroradiologist’s confidence in the resulting activity map. If the activity patterns of the patient are highly inconsistent across the run, the neuroradiologist might be able to retain some diagnostic information by re-analyzing those subsets of the data which are most indicative of task compliance. Splitting the data can also reveal if a patient’s activity pattern is atypical but stable, as was the case for our face condition. Here, we saw that although the patterns were not as expected, they were highly similar across blocks. Such analyses might allow to better decide if an inconclusive looking activity pattern is due to noise or is a veridical representation of an unexpected cognitive strategy the patient engaged in.

### Pattern analysis methods do not have to be ‘blackbox’

The pattern analyses in the present study were all based on the notion of minimizing the sums of squared differences between two observations. While these methods certainly do not take advantage of all the information contained in the data, they are highly versatile and robust (Hilborn and Mangel, 1997), and work well for fMRI data (Haxby, 2001). With the rise of artificial intelligence methods in medical imaging (Esteva et al., 2017), there is growing concern that the decisions made by algorithms might be excellent but the reasoning behind them will remain impenetrable to a human (Castelvecchi, 2016). Therefore, it could prove beneficial to accompany methods of high sophistication with more transparent (‘glass box’) analyses like the present ones.

### Localization and decoding of functions is complementary

While presurgical diagnostics are usually only concerned with brain mapping, the main benefit of decoding might be to better understand what exactly is being mapped. Even for a well-defined language task, the way the task is performed will not be identical for two different individuals. One patient might produce an activity pattern encompassing Broca’s area, SMA,Wernicke’s area and visual wordform area (VWFA). This patient’s activity would allow for a relatively safe interpretation of lateralization, depending on whether this network of activation is localized in the left or right hemisphere. If, on the other hand, another patient’s activity pattern for the language task resembled a default mode network, one would conclude that the task was not performed at all and not use the map to determine the degree of lateralization. Between these two extreme cases, a whole continuum of prototypical vs. improper task performance will occur in clinical practice. For example, a language production task will pose different demands on working memory or executive functions, depending on how difficult a patient finds the task overall, or for a specific category. The same frontal activity could be part of a strongly lateralized language network, or of a bilateral task-positive network, which typically includes frontal and parietal areas. Whether one is willing to use the frontal activity to draw conclusions about the language lateralization of a patient, could thus depend on how strongly each of those patterns is expressed in the whole-brain activity map. Therefore, the localization of functions (knowing where things are) could be aided by quantifying what cognitive demands the task poses for each patient (knowing what is being mapped). However, for such an approach to make sense, one has to derive maximally independent information for both localization and decoding. One possibility might be to use information at different spatial scales: Confidence in determining the dominant hemisphere would be conditional on how plausible the activity patterns within the hemispheres looked like. Another possibility might be to use different regions for localization and decoding: The confidence in a frontal activity corresponding to Broca’s area would depend on whether the region co-activates with other language-related areas in the rest of the brain. Finally, one could use different contrasts. For example, decoding would be performed only during rest, to check for prototypical activity in the default mode network. If this is established, an unconstrained analysis of activity during task performance would be performed. In the case of processing language, semantic maps of word meaning have been shown to be represented as very fine-grained and globally distributed activity patterns, comprising regions which are not part of the core language network (Huth et al., 2016). If it would be possible to decode the content of each block (e.g. producing names of animals vs. names of tools in a verbal fluency task), this would allow for a very close monitoring of the patient’s covert behavior, independent of the patient’s lateralization. Therefore, such types of decoding could be a valuable substitute for the lack of behavioral output which currently limits the applicability of mental imagery tasks.

### Conclusion

The present study showed how brief periods of covert thought can be decoded regarding the cognitive domains involved. The categories of language production, motor imagination, visuo-spatial navigation, face processing, and task-free resting were reliably differentiated using basic similarity metrics. This was possible using both training data from the same participant as well as independent meta-analytical data from other studies, which allow for immediate decoding without prior training. Capitalizing on the non-invasive nature of fMRI, we showed how exploratory approaches towards collecting and analyzing fMRI data can provide new impulses regarding its application in the individual case.

## Additional Information

### Acknowledgments

We would like to thank Anke Diekmann for help with data collection.

### Supporting Information

**S1. Instructions.** Reference guide for instructing a participant to perform the five mental imagery tasks.

**S2. Study Design.** Full study design with onsets and durations of all blocks.

### Copyright

(C)2018 Wegrzyn et al. This article is distributed under the terms of the Creative Commons Attribution License, which permits unrestricted use, distribution, and reproduction in any medium, provided the original authors are credited.

### Data Availability

Raw functional MRI data are available on openneuro.org/datasets/ds001419.

Preprocessed functional MRI data are available on doi.org/10.6084/m9.figshare.5951563.v1.

Results maps are available on neurovault.org/collections/3467

Code to reproduce the results and figures, as well as to recreate this manuscript, can be found on doi.org/10.5281/zenodo.1323665.

